# Regulating white blood cell activity through the novel Universal Receptive System

**DOI:** 10.1101/2025.01.06.631232

**Authors:** Victor Tetz, Kristina Kardava, Maria Vecherkovskaya, Alireza Khodadadi-Jamayran, Aristotelis Tsirigos, George Tetz

**Affiliations:** Human Microbiology Institute, New York, NY 10014, USA; Tetz Labs, New York, NY 10014, USA; Applied Bioinformatics Laboratories, NYU School of Medicine, New York, NY 10016, USA; Department of Pathology, NYU School of Medicine, New York, NY 10016; Department of Medicine, Division of Precision Medicine, NYU School of Medicine, New York, NY 10016

**Keywords:** antimicrobial, anticancer, leukocyte, activity, DNA-based receptors, RNA-based receptors, immune cells, Leukocyte-Tells, Teazeled receptors, TezR, phagocytosis, Universal Receptive System, New Biology, While blood cells

## Abstract

The understanding of the mechanisms that control key features of immune cells in various disease contexts remains limited, and few techniques are available for manipulating immune cells. Thus, discovering novel strategies for regulating immune cells is essential for gaining insight into their roles in health and disease. In this study, we investigated the potential of the recently described Universal Receptive System to regulate human immune cell functions. This was achieved for the first time by specifically targeting newly discovered surface-bound DNA and RNA-based receptors on leukocytes and generating “Leukocyte-Tells.” This approach upregulated numerous genes related to immune cell signaling, migration, endocytosis, and phagocytosis pathways. The antimicrobial and anticancer activities of Leukocyte-Tells exceeded the activity of control leukocytes *in vitro*. In some settings, such as in antibiofilm experiments, the Leukocyte-Tells showed up to 1,000,000-fold higher activities than control leukocytes. Our findings reveal, for the first time, that the Universal Receptive System can orchestrate fundamental properties of immune cells, including enhanced antimicrobial and anti-tumor activities. This novel approach offers a new avenue for understanding the biology and regulation of white blood cells.

## Introduction

Despite substantial progress in developing novel techniques aimed at understanding cellular regulation and activity, a considerable knowledge gap remains in this area of research [1]. Commonly used techniques related for modulating white blood cell (WBC) activity affecting cell activation, proliferation, differentiation, and genetic editing, are rooted in what we traditionally define as “Classical Biology” [2–5]. Although numerous cellular pathways have been identified and described, the exact mechanisms underlying their interplay, coordination, and adaptation to various stimuli remain elusive [6]. These knowledge gaps limit the targeted modulation of cellular characteristics ().

Our recent work describes, for the first time, a Universal Receptive System that is present in both prokaryotic and eukaryotic organisms. This system is composed of DNA- and RNA-based elements with receptive and regulatory functions, designated as Teaseled receptors (TezRs), located extracellularly, along with reverse transcriptases and recombinases, which are implicated in downstream signaling processes [7,8]. The discovery of DNA and RNA based TezRs as receptors and regulators, made by our team, was later supported by artificially constructed DNA receptors [9].

Previously, we demonstrated the effectiveness of targeting the Universal Receptive System to promote the emergence of novel cellular states and properties without directly affecting the genome [7,8,10,11]. Specifically, we observed the system’s primary role in cellular responses in prokaryotes and eukaryotes to various chemical, physical, and biological factors, as well as in controlling cell memory and forgetting. Moreover, we documented significant changes in the gene expression patterns of genes involved in essential cellular processes, including energy metabolism, proliferation, apoptosis, and protein expression [10].

Notably, our previous findings expanded the known role of the Universal Receptive System in controlling some of the fundamental properties of immune cells [12]. Specifically, we showed that the Universal Receptive System can control the production of diverse bioactive molecules (including antimicrobial and anticancer compounds) by leukocytes. Thus, our findings highlight the broad potential of targeted modulation using the Universal Receptive System to orchestrate diverse cellular processes. This approach substantially affects cellular activities and holds promise as a tool in “New Biology” for engineering novel cellular states. This finding highlights the novel roles of DNA and RNA, which are distinct from their traditional gene-encoding and gene-translating functions. This discovery is further supported by additional data revealing direct interactions between extracellular DNA and proteins, a phenomenon termed the “Pliers function.” These interactions induce altered protein folding and the generation of novel protein isoforms, resulting in changes of already synthesized extracellular proteins [13–15].

Leukocytes comprise a heterogeneous population of highly specialized nucleated immune cells. They actively engage in immune surveillance and execute immune responses, enabling eradication of infectious pathogens and cancers. Three major leukocyte subpopulations (lymphocytes, monocytes, and granulocytes), which are distinguished by their cellular architecture, granularity, and nuclear morphology, operate in distinct ways to contribute to immune protection [16][17]. While some leukocytes directly recognize and eliminate infected or cancer cells through cytolytic activity and cytokine release, others function indirectly by releasing danger signals, producing antimicrobial and anticancer peptides, mediating immune-cell recruitment, and serving as scavengers and antigen-presenting cells [18]. These highly regulated functions ensure control over diverse immunological threats and minimize the risk of undesired immune responses associated with various immunopathologies.

Importantly, relatively few methods exist for modulating the fundamental properties of leukocytes *ex vivo* to increase their antimicrobial or anticancer activities. Those methods are primarily based on stimulation with cytokines and different culture conditions [19,20]. For instance, the activity of T cells can be enhanced using a combination of anti-CD3 antibodies and a costimulatory signal (such as an anti-CD28 agonist) or natural killer cells stimulated with IL-2 and IL-15, as well as dendritic cells activated through Toll-like receptors interacting with pathogen-associated molecular patterns [21][22].

Gene editing is another strategy used to modify the anticancer effector function of the immune system, with Chimeric Antigen Receptor T (CAR-T) cells representing the most notable strategy [23,24]. This approach has demonstrated efficacy in treating several blood tumors, including B-cell leukemias, lymphomas, and multiple myelomas [25–27]. CAR-T cells are generated by modifying patient-derived T cells to express synthetic protein receptors, enabling them to eliminate target cells expressing the corresponding antigen [28]. Another approach utilized in experimental therapies involves the genetic modification of peripheral blood T cells *ex vivo* to express tumor-specific T cell receptors that target specific tumor antigens [29,30].

Although these methods have significantly advanced the field of immune cell programming and modulation, they have remained confined to a relatively restricted spectrum of options, emphasizing the need for alternative approaches to regulate the fundamental properties of leukocytes [31,32].

In this study, we discovered a critical and previously unknown role of the Universal Receptive System in orchestrating the key characteristics of human immune cells. Our findings demonstrate that deactivating the recently discovered TezRs led to the emergence of leukocytes (named here as “Leukocyte-Tells”) with upregulated antimicrobial and anticancer activities over the levels described for WBCs. This discovery marks a significant step forward in using a Universal Receptive System to orchestrate numerous critical pathways within immune cells, thereby enhancing our understanding of their biology in the context of various diseases.

## Results

### Leukocyte-Tells exhibited enhanced antimicrobial activity

Initially, we evaluated whether Leukocyte-Tells exhibited increased antimicrobial activity, ensuring that the process used to generate these cells did not affect their composition or viability (Supplementary Table S1).

We determined the minimum inhibitory concentration (MIC) and minimal fungicidal concentration (MFC), which represent the lowest concentrations of leukocytes required completely inhibit bacterial and fungal growth, respectively [33,34]. The MICs of Leukocyte-Tells and Leukocyte-Control cells against various bacterial and fungal strains are shown in Table 1. Notably, Leukocyte-Tells exhibited 10–10,000 higher activity against all tested bacterial and fungal strains than the controls (Table 1).

**Table 1.** Lowest dilutions of leukocytes required to completely inhibit visible bacterial and fungal growth.

Moreover, a clear dose-dependent effect of Leukocyte-Tells on bacterial growth was evident (Figure 1A), where higher Leukocyte-Tells: bacterium ratios correlated with lower bacterial growth.

**Figure 1:**
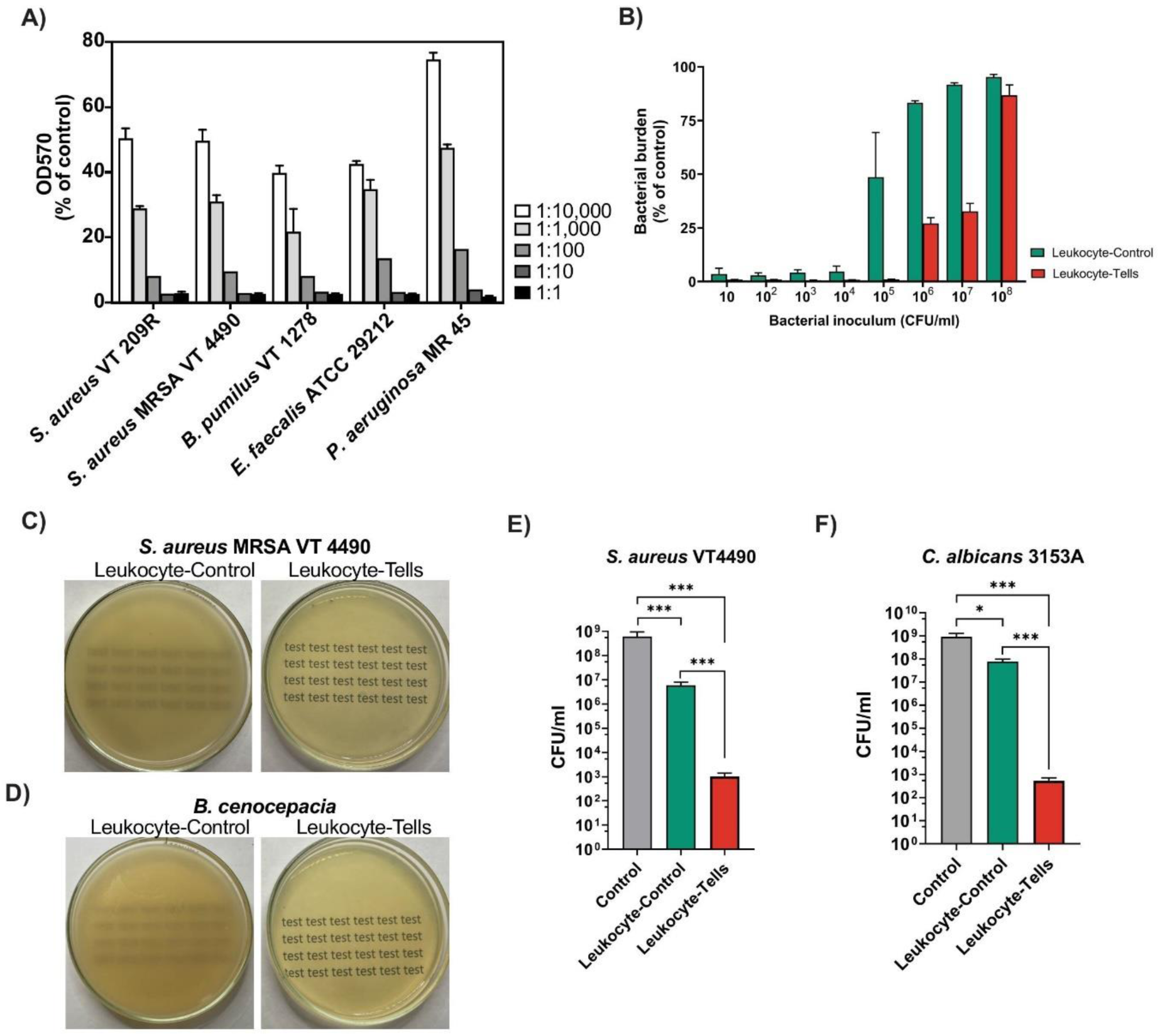
Antimicrobial activity of Leukocyte-Tells. (A) Quantitative analysis of the dose-dependent effect of Leukocyte-Tells on growth of various bacterial species. The optical density (OD) of each well was measured at 570 nm (OD570). The values are depicted as % of control, wells seeded with bacterial only, without leukocytes. Bar graphs represent mean±SD (n=3). Representative of 3 independent experiments. (B) Quantitative analysis of Leukocyte-Tells’ effect on increasing *S. aureus MRSA VT 4490* loads. Fixed number of Leukocyte-Tells (10^5 leukocytes/mL) was tested against increasing bacterial load. Bar graphs represent mean±SD (n=3). Representative of 3 independent experiments. Multiple unpaired t-tests. (C-D) Effect of Leukocyte-Control and Leukocyte-Tells on growth of *S. aureus MRSA VT 4490* (C) and *B. cenocepacia* (D) evaluated using agar overlay method. (E-F) Effect of Leukocyte-Control and Leukocyte-Tells on preformed biofilms of (E) *S. aureus VT 4490* and (F) *C. albicans* 3153A Bar graphs represent mean±SD (n=3). Representative of 3 independent experiments.

We further assessed the growth-inhibitory effect of increasing the bacterial load, using *Staphylococcus aureus* MRSA VT 4490 as an example, in the presence of 10^5^ Leukocyte-Tells or Leukocyte-Controls (Figure 1B). Notably, bacterial-growth inhibition with Leukocyte-Controls was observed only when the bacterial load was less than 10^5^ colony-forming units (CFUs). Starting with 10^5^ CFUs, bacterial growth reached approximately 50% of that in the control group. In contrast, Leukocyte-Tells substantially controlled bacterial growth, completely inhibiting bacteria at the initial *S. aureus* inoculum of 10^5^ CFUs and, even at higher initial inoculum of 10^7^ CFUs, reducing the bacterial load to approximately 30% of the control level.

Similarly, using the agar-overlay method, we demonstrated that Leukocyte-Tells had greater antimicrobial activity than Leukocyte-Controls, even when challenged with a high inoculum of gram-positive (*S. aureus*) (Figure 1C) or gram-negative bacteria (*B. cenocepacia*) (Figure 1D) [35].

Furthermore, we assessed the effect of Leukocyte-Tells on preformed biofilms, which are characterized by higher resistance to immune cells Immunometabolism in biofilm infection: lessons from cancer [36,37]. Leukocyte-Tells reduced the viable cell counts in 24 h old *S. aureus* biofilms by 100,000–1,000,000-fold than control treatment and were 100–10,000-fold more active than the Leukocyte-Controls (Figure 1E). Even more striking differences were observed in 24 h old *Candida albicans* biofilms, which were particularly insensitive to Leukocyte-Controls (Figure 1F). Under the same conditions, Leukocyte-Tells reduced the viable counts of fungal biofilms to 100,000–1,000,000-fold lower levels in untreated biofilms and were 10,000–1,000,000-fold more effective than Leukocyte-Controls.

Thus, Leukocyte-Tells exhibited greater antimicrobial activity than Leukocyte-Controls with various bacterial and fungal species, suggesting their superior potential to control high microbial burdens.

### Leukocyte-Tells exhibited enhanced in vitro anticancer activity

Based on the results of our antimicrobial-activity studies, we investigated whether Leukocyte-Tells might exhibit higher anticancer activity than Leukocyte-Controls. To this end, we analyzed the anticancer effects of Leukocyte-Tells and controls against a non-small cell lung cancer (NSCLC) cell line (NCI-H1299) and a glioblastoma cell line (T87G). Assessing tumor cell viabilities showed that the viabilities of the NCI-H1299 and T98G cells were significantly lower after treatment with Leukocyte-Tells than with the control-group cells (Figure 2A, B). Furthermore, NCI-H1299 monolayers treated with Leukocyte-Controls were uninterrupted and confluent, whereas those treated with Leukocyte-Tells exhibited extensive apoptosis, with numerous disrupted areas (Figure 2C). Thus, the destruction of TezRs in leukocytes enhanced their activation state, which improved their anticancer activity.

**Figure 2:**
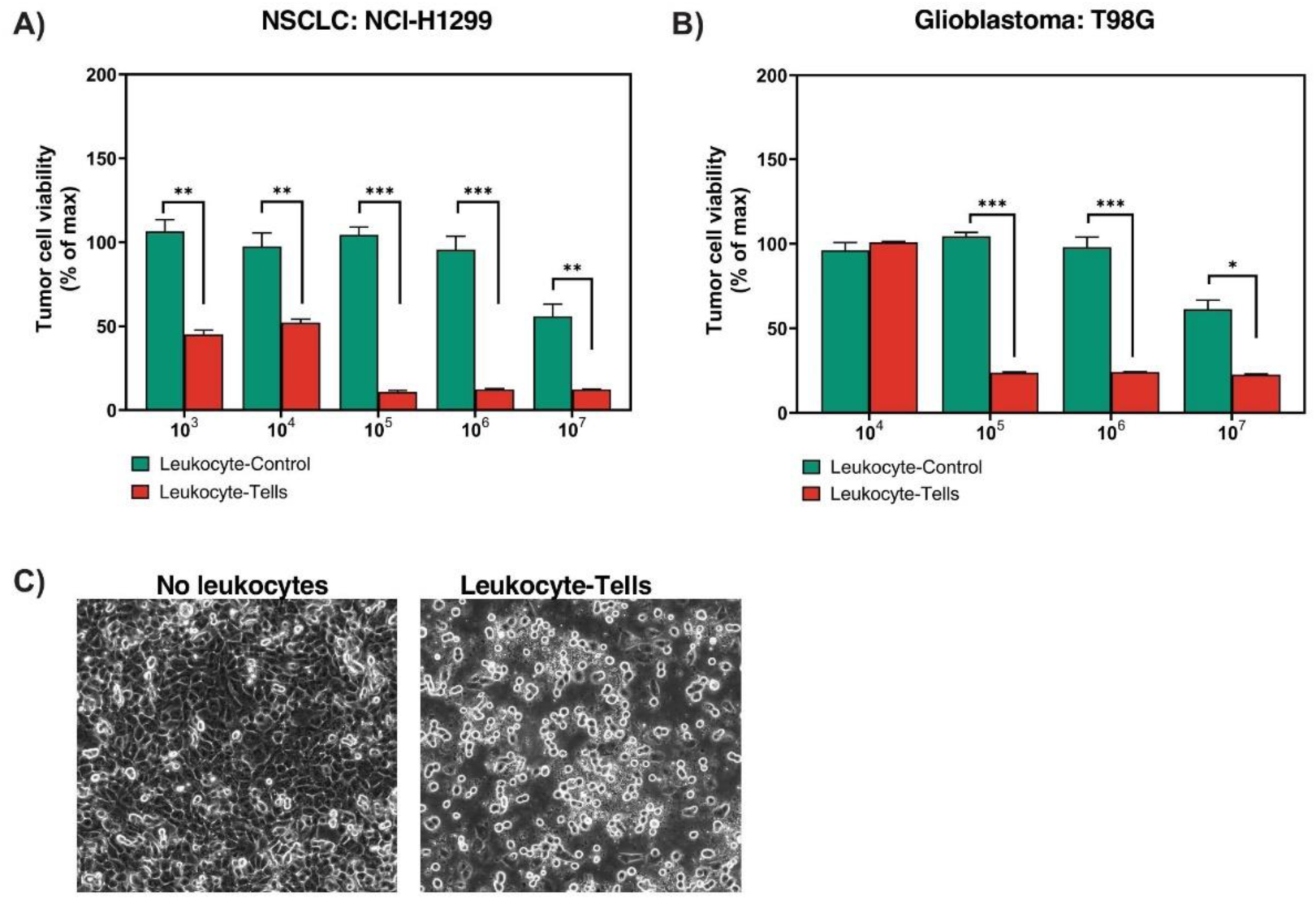
Anti-cancer activity of Leukocyte-Tells. Viability of human (A) NSCLC, NCI-H1299 and (B) human glioblastoma, T98G tumor cell lines in presence of increasing numbers of Leukocyte-Control or Leukocyte-Tells, expressed as % of the maximal viability measured in the absence of leukocytes. Bar graphs represent mean±SD (n=3). Two-way ANOVA. (C) Growth of NSCLC, NCI-H1299 tumor cell line in culture in presence of Leukocyte-Tells.

To support our initial findings demonstrating the superior antimicrobial and anticancer properties of Leukocyte-Tells, we conducted genome-wide mRNA-expression profiling and functional pathway analysis with both Leukocyte-Tells and untreated control leukocytes (Figure 3A–H). Our analysis revealed 1,496 differentially expressed genes (DEGs) between the Leukocyte-Tells and Leukocyte-Control cells. Most DEGs (1,281 of 1,496) were upregulated in Leukocyte-Tells, although some DEGs (215 of 1,496) were downregulated (Figure 3A).

**Figure 3:**
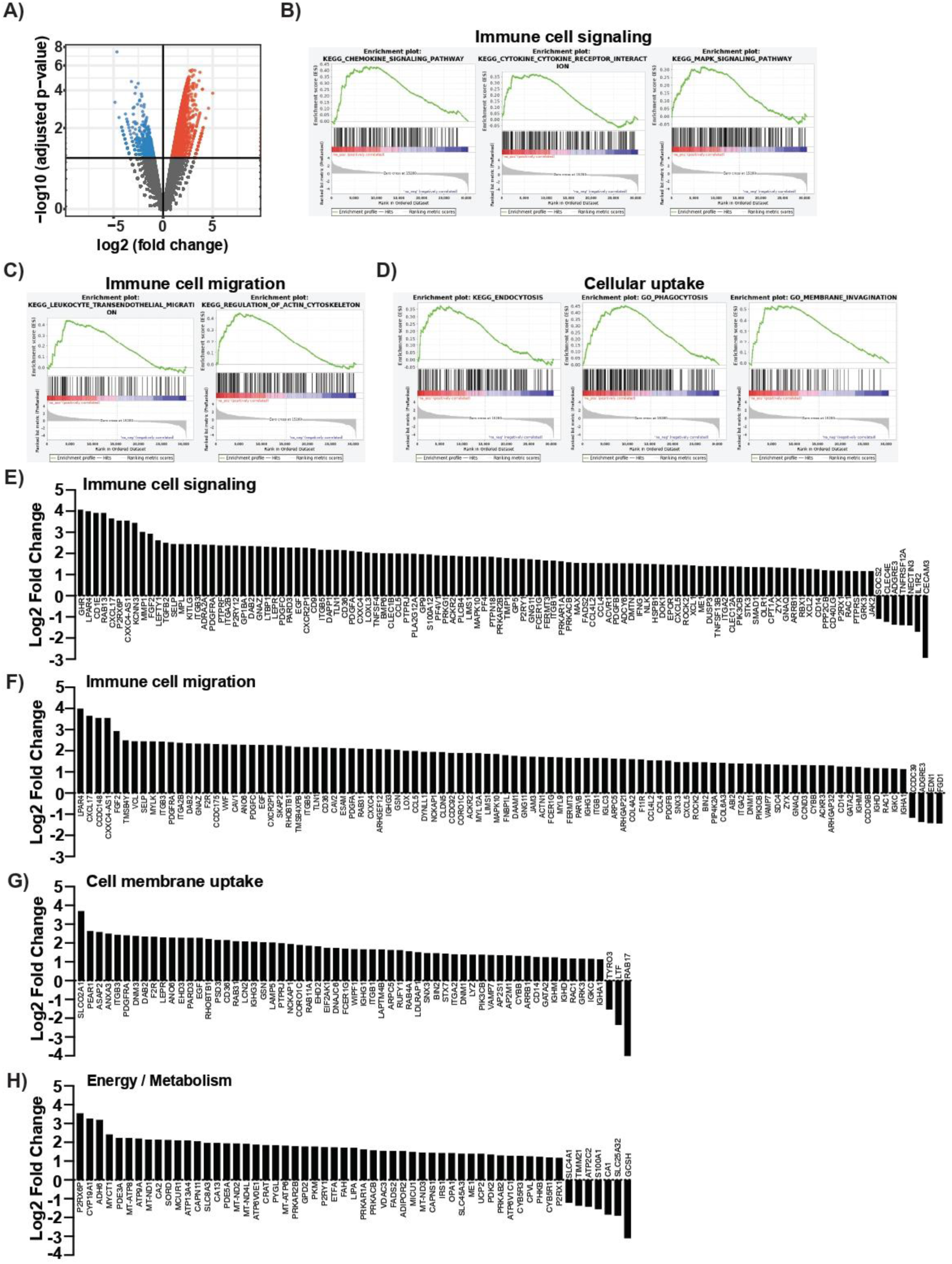
Gene expression analysis of Leukocyte-Tells. (A) Genes differentially expressed in Leukocyte-Tells versus Leukocyte-Control. (B-D) Gene set enrichment analysis plots of selected functional pathways are presented for Leukocyte-Tells vs. Leukocyte-Control. Functional pathways are grouped in three categories: (B) Immune cell signaling, (C) Immune cell migration and (D) Cell membrane uptake. (E-F) Differentially expressed genes associated with four distinct functional pathways.

To further contextualize the gene-expression changes that occurred in Leukocyte-Tells, we performed gene set enrichment analysis [38,39]. Our analysis revealed enrichment for genes associated with numerous functional pathways in Leukocyte-Tells (Figure 3B–D). The identified enriched pathways were grouped into three distinct and broad categories: immune cell signaling (Figure 3B), immune cell migration (Figure 3C), and cell membrane uptake (Figure 3D). Within the cell signaling group, key pathways enriched in Leukocyte-Tells included chemokine signaling, cytokine signaling, and mitogen-activated protein kinase signaling, all of which are indicative of increased Leukocyte-Tell activity [40–42]. Similarly, the enrichment for pathways, such as leukocyte transendothelial migration and actin cytoskeleton remodeling were suggested that the mobility of the immune cells had increased, which is characteristic of acute inflammation-associated responses (Figure 3C) [43–45]. Finally, enrichment in terms of endocytic and phagocytic pathways indicated that enhanced cell membrane uptake occurred, representing another key feature of different activated immune cell subtypes (Figure 3D) [46,47].

Next, we analyzed just the DEGs related to the same three functional pathway categories (Figure 3E–G). The most significantly upregulated genes in leukocytes within the immune cell-signaling pathway included numerous cytokines and cytokine receptors, such as CXCL17, CCR3, CCL4, and XCL2 (Figure 3E) [48–50]. Additionally, leukocytes were characterized by significant upregulation of GHR and PTPRF, which help initiate immune cell activation and regulate various immune-signaling pathways through the phosphorylation of target proteins, respectively (Figure 3E) [51,52]. Some of the most highly upregulated genes in leukocytes during immune cell migration were ESAM, JAM3, RAP1GDS1, and RAPGEF5 (Figure 3F) [53–56]. Finally, in terms of immune cell uptake, the most upregulated genes were associated with endocytosis, vesicle trafficking, and cytoskeleton reorganization, such as SLCO2A1, ANXA3, DNM3, and EHD3 (Figure 3G) [57–60]. We also observed substantial alterations in energy-associated pathways in Leukocyte-Tells, marked by the upregulation of genes implicated in metabolism, such as GPD2, FAH, PYGL, and PDK2 (Figure 3H) [61–64]. These genes are associated with alternative energy supply pathways, notably glycolysis [64].

We identified a substantial proportion of non-coding RNAs (ncRNAs) among the DEGs, including long intergenic non-protein-coding RNAs and pseudogenes (Supplementary Tables S2 and S3). These elements function as master regulators of gene expression and play crucial roles in orchestrating complex cellular processes, such as apoptosis, growth, differentiation, and proliferation [65–67]. Interestingly, these ncRNAs and pseudogenes were the most significantly upregulated and downregulated DEGs in Leukocyte-Tells, respectively.

Thus, the Leukocyte-Tells showed gene-expression patterns suggestive of an enhanced activation state [68,69].

### Leukocyte-Tells exhibited enhanced cytokine production upon stimulation

Cytokines play crucial roles in orchestrating diverse cellular processes within the immune system, including the proliferation, migration, and differentiation of immune cells [70–72]. Here, we compared the cytokine-production profile of Leukocyte-Tells after lipopolysaccharide (LPS) stimulation with that of Leukocyte-Control cells (Figure 4).

**Figure 4:**
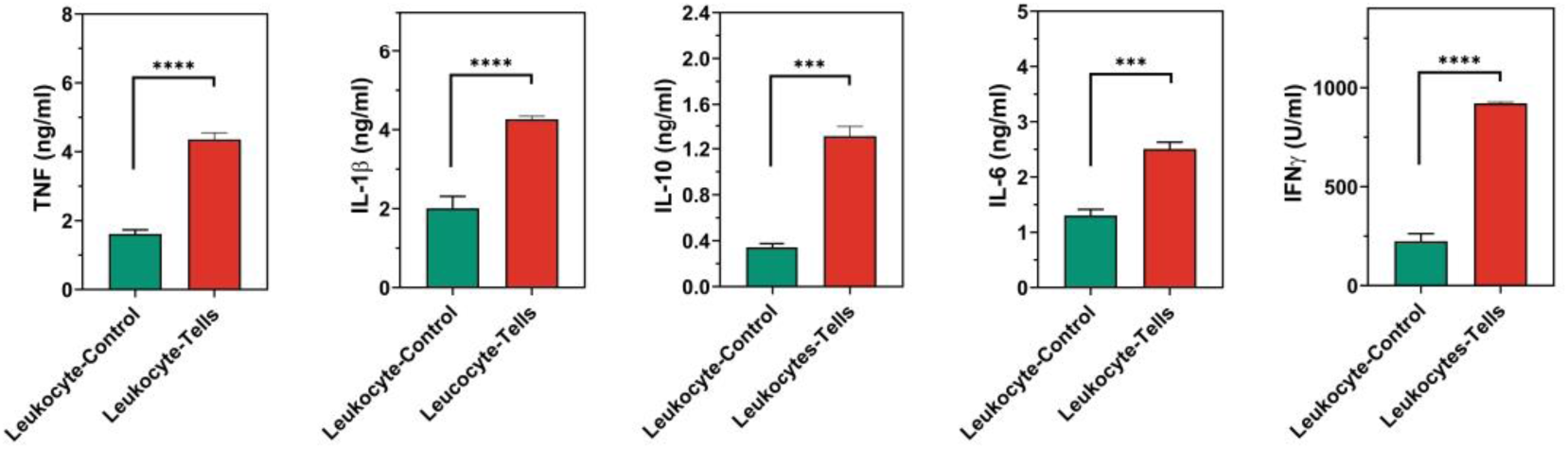
Cytokine production of LPS-stimulated Leukocyte-Tells. Concentration of TNF, IL-1*β*, IL-6, IL-10 and IFN-*γ* (from left to right) measured in supernatants of Leukocyte-Control and Leukocyte-Tells cultures after one hour of LPS stimulation. Bar graphs represent mean±SD (n=3). Representative of 3 independent experiments. One-way ANOVA.

We focused on five cytokines, namely TNF, IL-1β, IFN-γ, IL-6, and IL-10, which have pleiotropic and context-dependent roles [73–76]. Notably, the Leukocyte-Tells showed significantly higher levels of all cytokines studied after LPS stimulation than Leukocyte-Control cells (Figure 4). Consistent with the gene-expression data indicative of an activated signature in Leukocyte-Tells, these cells exhibited greater expression of key effector cytokines in response to stimulation.

### Leukocyte-Tells exhibited greater phagocytic activity and production of granule enzymes

Gene-expression analysis clearly indicated that the Leukocyte-Tells showed enhanced activity and augmented cellular-uptake processes, such as phagocytosis (Figure 5). To formally assess the phagocytic activity of leukocytes, we compared their ability to phagocytose *Escherichia coli* with that of Leukocyte-Controls. Our findings confirmed the initial gene-expression analysis, revealing that Leukocyte-Tells showed approximately 20% higher bacterial internalization than their Leukocyte-Control counterparts (Figure 5).

**Figure 5:**
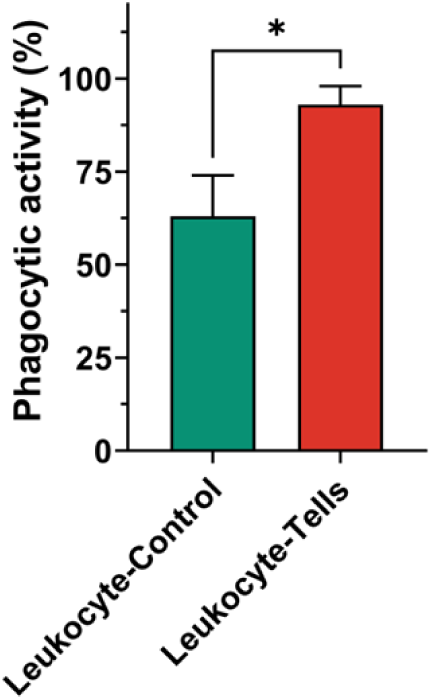
Phagocytosis of Leukocyte-Tells. Phagocytic activity of Leukocyte-Tells were compared with Leukocyte-Control Bar graphs represent mean±SD (n=3). Representative of 3 independent experiments. One-way ANOVA.

## Discussion

Previously, we demonstrated the importance of the Universal Receptive System in regulating diverse cellular functions, including the production of antimicrobial and anticancer bioactive compounds (AABCs) by leukocytes [7,12]. In this study, we evaluated its impact on shaping crucial features of human immune cells.

By leveraging the Universal Receptive System, we generated Leukocyte-Tells with significantly higher antimicrobial and anticancer activities than the original immune cells. This was accomplished by selectively inactivating extracellular DNA- and RNA-based TezRs receptors. Depleting the TezRs led to the concurrent modulation of multiple pathways implicated in leukocyte immune responses, with substantial changes in gene-expression patterns and upregulation of key pathways associated with immune cell function, including migration, immune cell signaling, the use of alternative energy sources, and cell membrane transport.

In this study, we utilized leukocytes from multiple donors, which revealed consistent activation and highly reproducible patterns across all tested samples. To emphasize the reproducibility of the cell responses controlled by the Universal Receptive System, we suggest using the term “Cell Genome Memory Managing” to indicate that the induced cellular states are pre-programmed and can be reactivated.

The results of our *in vitro* antimicrobial studies represent a significant advancement, revealing a stark difference in potency between Leukocyte-Tells and their control counterparts, which were isolated from the same multiple donor group. Specifically, the Leukocyte-Tells demonstrated a striking increase in potency (up to 100,000-fold) against bacteria and fungi. Moreover, Leukocyte-Tells exhibited stronger antibiofilm activity against microbial communities, which are known to be resilient against immune cells [37]. Furthermore, the superior antimicrobial activity of Leukocyte-Tells persisted irrespective of microbial resistance to antibiotics, highlighting the robustness of immune cell-based therapies in overcoming bacterial resistance.

In addition to their enhanced antimicrobial activity, Leukocyte-Tells showed superior anticancer activity *in vitro*, as evidenced by a >10,000-fold increase in efficacy against a p53-null NSCLC tumor cell line and a 100-fold increase in efficacy against a p53-mutant glioblastoma cell line (relative to those of control cells) [77,78].

The question arises as to whether the observed changes were directly associated with the activity of the Leukocyte-Tells themselves or indirectly caused by the increased secretion of AABCs in response to stimulation, as observed previously [12]. The antimicrobial and anticancer activities of the isolated AABCs from Leukocyte-Tells, when applied in our lab to the same microbial strains and cancer cell lines without Leukocyte-Tells, were lower than those shown by the Leukocyte-Tells, the observed effects were not solely attributable to AABCs [12]. These results indicated that other types of antimicrobial and anticancer effects generated by the Leukocyte-Tells also contribute to the observed effects.

The pronounced antimicrobial and anticancer activities of Leukocyte-Tells may also be partially attributable to increased phagocytosis. Furthermore, transcriptomic analysis confirmed that key genes associated with phagocytosis were induced, including members of the Rho family of GTPases and phosphatases [79]. Although the observed increase in phagocytosis was statistically significant, its magnitude was highly unlikely to solely account for the multi-fold increase in the antimicrobial and anticancer efficacies of Leukocyte-Tells versus those of the controls. Moreover, the Leukocyte-Tells exhibited increased production of cytokines, including TNF, IL-1β, IFN-γ, and IL-6. This elevation in cytokine release was associated with the activation, proliferation, and enhanced cytotoxicity of leukocytes and the production of reactive oxygen and nitrogen species [72]. We plan on continuing to perform an in-depth study of the antimicrobial and anticancer activities of Leukocyte-Tells in relevant animal models.

Interestingly, based on the transcriptomic profile, after the loss of TezRs in leukocytes, we observed the same pattern of upregulation of genes related to energy metabolism and migration as previously observed in prokaryotes and other eukaryotes [8]. For example, certain bacterial species show increased dispersion over unexpectedly long distances in response to TezR inactivation [10]. Similarly, we observed that multiple genes associated with migration and vascular extravasation were upregulated in Leukocyte-Tells. Furthermore, our previous findings revealed that modifying TezRs facilitated the growth of certain obligate aerobic bacteria in an anaerobic environment by upregulating glycolytic pathways, or by activating glycolytic pathways in bacteria cultivated in an aerobic environment following the loss of TezRs [8]. The latter observation agrees with the findings reported in this study, demonstrating the upregulation of multiple genes associated with accelerated metabolism and the utilization of alternative energy sources, including those involved in glycolysis (ADH and PKM) [80].

Importantly, in this study, we used leukocytes from multiple donors, which revealed consistent activation and a highly reproducible pattern across all tested samples. These data imply that the observed increase in anticancer and antimicrobial activities, although unusually high, was inherent to immune cells, as we did not modify the cellular genome; all modulations were performed by targeting TezRs.

In conclusion, our findings demonstrate that the distinctive attributes of Leukocyte-Tells hold strong potential for advancing the understanding of the full spectrum of roles played by the Universal Receptive System within cells. These observations warrant further evaluation of Leukocyte-Tells as antimicrobial and anticancer agents.

## Materials and methods

### Bacterial and fungal strains and culture conditions

*S. aureus* ATCC 29213, *Enterococcus faecalis* ATCC 29212, *E. coli* ATCC 25922, and *Mycobacterium smegmatis* ATCC 607 were purchased from the American Type Culture Collection (ATCC; Manassas, VA, USA). Clinical isolates of *S. aureus* VT 209R, *S. aureus* MRSA VT 4490, *Bacillus pumilus* VT 1278, *Klebsiella pneumoniae* VT 1698, and *Aspergillus niger* VT 1591 were obtained from a private collection (Human Microbiology Institute, NY, USA). *Pseudomonas aeruginosa* MR45, isolated from a patient with cystic fibrosis, was obtained from the CF Foundation Therapeutics Development Network Resource Center for Microbiology at Seattle Children’s Hospital (Seattle, WA, USA). *C. albicans* 3153A was provided by Dr. Paul Fidel (LSU Health).

Bacterial strains were passaged weekly on Columbia agar (BD Biosciences, Franklin Lakes, NJ, USA) and stored at 4 °C. All liquid subcultures were derived from colonies isolated from these plates and were grown in Mueller Hilton broth (Sigma-Aldrich, St. Louis, MO, USA), unless stated otherwise. For experiments involving solid medium, the bacteria were cultured on Columbia agar (Sigma-Aldrich). All cultures were incubated aerobically at 37 °C.

All fungal strains were subcultured from freezer stocks on Sabouraud dextrose agar (Oxoid, UK) plates and incubated at 30 °C overnight to obtain yeast and mold colonies for subsequent liquid subcultures on Sabouraud dextrose agar (Oxoid, UK).

### Cell lines

H1299 and T98G cell lines (ATCC) were authenticated via short tandem repeat profiling at our institute. For those experiments, the cells were cultured in RPMI-1640 medium (Sigma-Aldrich), with or without 10% fetal bovine serum (Gibco, Invitrogen Corporation, NY, USA), 1 mM L-glutamine (Sigma-Aldrich), penicillin G (100 U/mL), and streptomycin (100 mg/mL, Sigma-Aldrich), and were grown in a humidified chamber (95% air, 5% CO_2_) at 37 °C. The cells were detached from their flasks using trypsin/ethylenediaminetetraacetic acid (0.05%, Invitrogen).

### Generating Leukocyte-Tells

Donors were preselected who did not take any antibiotics, corticosteroids, or anticancer drugs in the last 3 months. Red blood cells (RBCs) were depleted from the blood of healthy donors by double-gradient centrifugation, as described previously, after obtaining informed consent. To generate Leukocyte-Tells after RBC removal, the remaining cellular components (including leukocytes and platelets isolated from several donors) were treated with DNase I and RNase A (1 µg/ml; Sigma-Aldrich), as previously described [7,12]. The control leukocytes (Leukocyte-Control cells) represented untreated leukocytes and platelets. We used five sets of Leukocyte-Tells from random donors of varied sex, age, and race in this study. All sets were processed without freezing within 24 h of generation. The number of leukocytes in the samples was determined using an automated SYSMEX XN-330 hematology analyzer (Sysmex, Japan). Cell viability was assessed using a Nucleocounter NC202 (Chemometech), with a viability of >70% required for use in this study.

### Total RNA extraction, purification, library preparation, and sequencing

The cells were harvested via centrifugation at 300 × *g* for 15 min and resuspended in phosphate-buffered saline (PBS). RNA was extracted from leukocytes using the RNeasy Mini Kit (Qiagen, Germantown, Maryland, USA). RNA quality was assessed spectrophotometrically at 230, 260, and 280 nm with a Promega GloMax Discover Microplate Reader (Promega, Madison, WI, USA). Ribodepletion was performed using the Ribo-Zero Magnetic Gold Kit (Epicenter, Madison, Wisconsin, USA). Transcriptome sequencing libraries were prepared from RNA using the TruSeq Stranded Total RNA Library Prep Kit and sequenced on an Illumina NovaSeq 6000 (Illumina, San Diego, California, USA) using a 2 × 150 nucleotide paired-end strategy (130 MM reads, maximum).

### RNA-seq data processing

All reads from the sequencing experiment were mapped to the reference human genome (GRCh38) using STAR (v2) Duplicate reads were removed using Picard tools (v1.126) (http://broadinstitute.github.io/picard/). Reads-per-million-normalized BigWig files were generated using BEDTools (v2.17.0) and bedGraphToBigWig tool (v4) [81]. Read count tables were generated using HTSeq (v0.6.0) and normalized based on library size factors using DEseq2, after which differential gene-expression analysis was performed. Transcripts with an adjusted p value of <0.05 and a log2 fold-change value of ±1.0 (2-fold) were considered significantly DEGs. All downstream statistical analyses and plot generations were performed using R (v3.1.1) (https://www.r-project.org/).

### Estimating MICs and MFCs in vitro

The MICs for bacteria and MFCs for leukocytes were determined using the serial microdilution method in accordance with a modified version of the Clinical and Laboratory Standard Institute guidelines [82]. The standard inocula for testing bacteria, yeast, and mold cells were 5 × 10^5^, 2.5 × 10^3^, and 5 × 10^4^ CFUs/mL, respectively. A 20 μL aliquot of the bacterial inoculum was added to the wells of 96-well plates containing 180 μL of leukocytes serially diluted (10-fold) in RPMI-1640 medium. Plates with bacterial cultures were incubated at 37 °C for 24 h, and plates with fungal cultures were incubated at 30 °C for 24 h. Untreated leukocytes from the same set of multiple donors were used as a negative control (Leukocyte-Controls). MICs and MFCs were defined as the lowest dilutions of leukocytes required to completely inhibit visible bacterial and fungal growth, respectively [83]. To evaluate microbial viability, bacterial and fungal suspensions were serially diluted, and 100 μL of diluted suspension was spread onto agar plates. Columbia agar plates were used for bacterial cultivation, and Sabouraud dextrose agar was used for fungal growth. Plates were incubated at 37 °C overnight, and CFUs were counted on the next day.

### Effect of leukocytes on bacterial biomass at planktonic growth

We inoculated wells of 96-well plates containing 180 µL of leukocytes (1 × 10^5^ cells/mL in RPMI-1640) with 20 µL of bacteria (from 10 to 1 × 10^8^ bacteria/mL). The plates were incubated at 37 °C for 24 h and the optical density (OD) of the microorganisms was evaluated using a Promega GloMax Discover Microplate Reader. The values were subtracted from the mean OD_570_ of wells containing 1 × 10^5^ leukocytes/mL.

### Agar-preincubation method

Petri dishes were filled with 20 mL 1.5% agar. The pre-autoclaved soft 0.7% agar overlay was melted at 60 °C in a water bath and allowed to cool without solidification. One hundred microliters of the tester strain culture was taken at 10^7^ bacteria/mL and tested leukocytes 10^7^ cells/mL) were tested on soft agar, vortexed to mix, and poured onto the surface of the 1.5% agar [84,85]. The plates were incubated at 37 °C for 24 h before the bacterial growth was assessed.

### Effect of leukocytes on bacterial and fungal biofilms

We added 200 µL *S. aureus* VT 4490 or *C. albicans* 3153A (each taken at 5 × 10^5^ CFUs/mL) in Mueller Hilton (Sigma) or Sabouraud dextrose broth (Oxoid), respectively, to the wells of 96-well flat-bottom microtiter plates [33]. Following a 24 h incubation at 37 °C, the resulting biofilms were washed twice with PBS to remove non-adherent bacteria, after which 200 µL leukocytes (1 × 10^5^ cells/mL) in RPMI-1640 was added. The plates were incubated at 37 °C for 24 h. After exposure, the contents of the wells were aspirated, and each well was washed three times with PBS. The biofilms were scraped thoroughly, with particular attention being paid to the edges of the wells [86]. The well contents were aspirated and the total CFU numbers were determined by serial dilution and plating bacteria and fungi on Columbia agar and Sabouraud dextrose agar, respectively.

### Tumor cell viability assay

Cell viability was determined by performing 3-(4,5-dimethylthiazol-2-yl)-2,5-diphenyltetrazolium bromide (MTT) assays. Briefly, H1299 or T98G cells (5 × 10^3^ cells/well) were seeded into 96-well plates in RPMI-1640 medium supplemented with 10% fetal bovine serum (FBS; Gibco) without antibiotics. After 24 h of incubation, the growth medium was replaced with fresh medium without FBS but with increasing concentrations of leukocytes (10^3^–10^8^ cells/ml). The control wells contained increasing concentrations of untreated leukocytes. The plates were incubated for 24 h. Each well was supplemented with 20 μL MTT solution (5 mg/mL in PBS) (Thermo Fisher, Waltham, MA), followed by additional 3 h incubation at 37 °C. Subsequently, the medium containing MTT was removed, and 50 μL of dimethyl sulfoxide (Sigma, USA) was added to each well to dissolve the formazan crystals. To assess the percentage of live cells in the samples, the absorbance was measured at 570 nm, while accounting for the background absorbance of leukocytes and cancer cells. The same method was used to evaluate the leukocyte viability after heating or protease treatment.

### Lactate dehydrogenase (LDH)-release assay

Cell viability was assessed using the CyQUANT LDH Cytotoxicity Assay (Thermo Fisher). Briefly, H1299 and T98G cells (5 × 10^3^ cells/well) were seeded in 96-well plates in RPMI-1640 medium. After a 24 h incubation, the growth medium was replaced with fresh medium containing increasing concentrations of leukocytes (10^3^–10^8^ cells/mL). Control wells contained increasing concentrations of leukocytes. The plates were incubated for an additional 24 h, and LDH release in each well was measured by measuring the absorbance at 490 nm. Cytotoxicity was determined by comparing LDH release to that of the positive control (cells treated with 1% Triton X-100) after subtracting the background values of leukocytes alone (not cocultured with cancer cells).

### Phagocytosis assays

The phagocytic activity of leukocytes was assessed using *E. coli* cultures. WBCs were collected as previously described and seeded in black-walled 96-well plates for 2 h at a density of 5 × 10^4^ cells/well. The *E. coli* cells were stained with fluorescein isothiocyanate (FITC; Thermo Fisher) and washed three times to remove the unbound dye. Phagocytosis was assessed by subtracting the average fluorescence intensity of the ‘no-cell’ negative controls from the experimental wells. We incubated 1 × 10^4^ leukocytes with Dextran-FITC (1 mg/ml) at 37 °C for 60 min, counter-stained the cells with an F4/80 antibody, and analyzed the cells using a BD FACSCanto flow cytometer (Becton Dickinson).

### Cytokine detection

To evaluate cytokine secretion in response to LPS, 10^5^ leukocytes were incubated with LPS (5 ng/mL) for 1 h. The concentrations of TNF, IFN-*γ*, IL-1*β*, IL-6, and IL-10 were measured in the culture medium using commercially available enzyme-linked immunosorbent assays (FineTest, Wuhan Fine Biotech).

### Statistical analysis

At least three biological replicates were studied under each experimental condition in three independent experiments using material from at least five donors, unless otherwise stated. Data are presented as the mean value ± standard deviation. Microbial quantification data were log_10_-transformed before analysis. The specific statistical tests used are indicated in the corresponding figure legends. GraphPad Prism version 10 (GraphPad Software, San Diego, CA, USA) or Excel 10 (Microsoft, Redmond, WA, USA) were used for statistical analyses and illustrations.

## Acknowledgements

Applied Bioinformatics Laboratories (ABL) is a shared resource partially supported by the Cancer Center Support Grant P30CA016087 at the Laura and Isaac Perlmutter Cancer Center. This work has used computing resources at the NYU School of Medicine High Performance Computing (HPC) Facility.

We would also like to thank Natalija Budimir for working on the manuscript.

## Data availability

Data availability is available upon request from the corresponding author.

---

**Supplementary Table S1.** Cellular contents of the Leucocyte-Control and Leukocyte-Tells populations

**Supplementary Table S2:** Non-protein-coding RNA in DEGs

**Supplementary Table S3:** Pseudogenes in DEGs

