## Supplemental Table 1 for "Regulating white blood cell activity through the novel Universal Receptive System"

**Supplementary Table S1. Cellular contents of the Leucocyte-Control and Leukocyte-Tells populations**

| **Cell type** | **Granulocytes (%)** | **Agranulocytes (%)** | **P value** |
| --- | --- | --- | --- |
| Leukocyte-Control | 60.7 | 39.3 | 0.395 |
| Leukocyte-Tells | 64.2 | 35.8 | 0.304 |

**Supplementary Table S2**

**Supplementary Table S3**
