## Supplemental Table 2 for "Regulating white blood cell activity through the novel Universal Receptive System"

**Supplementary Table S2: non-protein- coding RNA in DEGs**

AC000068.2

AC002070.1

AC002076.2

AC002381.1

AC002553.2

AC003682.1

AC004053.1

AC004584.1

AC004687.1

AC004840.1

AC005050.3

AC005332.6

AC005394.2

AC005410.1

AC005840.2

AC006116.1

AC007262.2

AC007383.2

AC007391.2

AC007431.3

AC007435.2

AC007998.3

AC008695.2

AC008763.1

AC008957.1

AC009032.1

AC009090.5

AC009127.1

AC009145.1

AC009159.2

AC009971.1

AC010186.1

AC010335.1

AC010737.1

AC011005.1

AC012313.10

AC012640.4

AC015849.1

AC015849.6

AC017002.5

AC017074.1

AC020558.2

AC020634.2

AC020658.2

AC020763.1

AC020910.4

AC020913.2

AC021054.1

AC021945.1

AC022126.1

AC022149.1

AC023024.1

AC023024.2

AC023632.5

AC023794.4

AC024940.2

AC025470.2

AC026117.1

AC026462.1

AC026951.1

AC027281.2

AC034102.3

AC037198.1

AC048338.2

AC055876.4

AC063944.2

AC064805.1

AC067852.5

AC067945.2

AC068234.2

AC068418.2

AC068759.1

AC069271.1

AC069368.1

AC073140.1

AC073332.1

AC073343.1

AC073842.1

AC073916.1

AC074351.1

AC078899.1

AC079296.1

AC079753.1

AC079753.2

AC079804.3

AC083870.1

AC084824.4

AC087071.2

AC087482.1

AC087878.1

AC090409.1

AC090409.2

AC090615.1

AC090772.3

AC090826.2

AC090844.2

AC090971.5

AC091057.1

AC091114.1

AC091173.1

AC091982.1

AC092135.3

AC092140.2

AC092535.2

AC092542.1

AC092723.5

AC092794.1

AC092919.2

AC093388.1

AC093512.2

AC093627.6

AC093909.1

AC095033.1

AC096564.2

AC096577.1

AC096638.2

AC096642.1

AC097460.3

AC097709.1

AC098614.3

AC098850.3

AC100863.1

AC104083.1

AC104794.3

AC104850.2

AC104971.2

AC105020.2

AC105415.1

AC106874.1

AC107027.1

AC107223.1

AC108136.1

AC109583.2

AC110611.2

AC112184.1

AC112198.2

AC112253.1

AC112254.1

AC113143.1

AC113361.1

AC113386.1

AC113404.1

AC113404.3

AC114495.2

AC114752.2

AC116903.2

AC117422.1

AC120114.1

AC120498.9

AC123912.4

AC124016.1

AC125232.2

AC127070.3

AC127502.2

AC134878.2

AC135584.1

AC136297.1

AC136628.3

AC138649.2

AC138761.1

AC139792.2

AC140125.2

AC144521.1

AC145285.3

AC147651.1

AC147651.4

AC211469.1

AC231533.1

AC241377.4

AC243960.2

AC246787.2

AL021707.9

AL021807.1

AL022324.2

AL022344.1

AL022345.4

AL023755.1

AL031289.1

AL031777.1

AL031963.3

AL033543.1

AL034397.3

AL035413.2

AL049610.1

AL078645.1

AL080250.1

AL117336.1

AL118508.3

AL118558.4

AL121601.2

AL121899.1

AL121899.4

AL133243.1

AL133444.1

AL133517.1

AL135903.1

AL137013.1

AL137847.2

AL138689.2

AL138713.1

AL138756.1

AL139089.1

AL139095.4

AL157834.2

AL157895.1

AL158206.1

AL158211.3

AL160396.1

AL162424.1

AL163932.1

AL354718.1

AL355073.1

AL356414.1

AL356475.1

AL359643.2

AL359644.1

AL359880.1

AL360015.1

AL360227.1

AL390728.6

AL390755.3

AL390882.1

AL391001.1

AL391422.4

AL391903.1

AL442067.2

AL445224.1

AL445426.1

AL450267.1

AL450267.2

AL451050.2

AL451164.1

AL499627.1

AL512652.2

AL591135.1

AL591506.1

AL603840.1

AL627171.1

AL731557.1

AL731577.2

AP000547.3

AP000692.2

AP000866.2

AP001029.2

AP001033.1

AP001033.2

AP001102.1

AP001189.1

AP001189.3

AP001267.5

AP001610.2

AP001636.2

AP001636.3

AP001992.2

AP002449.1

AP002478.1

AP002505.3

AP002992.1

AP003049.2

AP003068.2

AP003080.1

AP003392.4

AP003469.1

AP005482.3

LINC00211

LINC00355

LINC00504

LINC00520

LINC00534

LINC00635

LINC00664

LINC00674

LINC00853

LINC00892

LINC00958

LINC00989

LINC01011

LINC01088

LINC01151

LINC01182

LINC01341

LINC01476

LINC01618

LINC01750

LINC01923

LINC01992

LINC02150

LINC02211

LINC02280

LINC02284

LINC02291

LINC02340

LINC02384

LINC02470

LINC02539

LINC02668

LINC02715

LINC02770

LINC02842
