## Supplemental Table 3 for "Regulating white blood cell activity through the novel Universal Receptive System"

Supplementary Table S3: Pseudogenes in DEGs

ACTBP11

ACTBP2

ACTBP7

ACTG1P14

AGGF1P10

BIN2P1

BUD13P1

CAP1P2

COX6CP18

CXCR2P1

E2F6P4

EEF1A1P24

EIF4EP2

ELL2P1

FTH1P11

FTH1P2

FTH1P20

FTH1P23

FTH1P8

FTLP2

FTLP3

GAPDHP1

GCNT1P1

GLYATL1P1

GLYATL1P2

GNAQP1

GPX1P1

GPX1P2

H3F3AP4

H3F3AP6

HNRNPA1P15

HNRNPA3P15

HSPA8P3

HTATIP2

KLF3P1

KRT8P39

LAMP5

LARP1P1

MAP2K4P1

MEIS3P2

MTATP6P1

MTCO1P12

MTCO1P2

MTCO2P12

MTCO3P12

MTCYBP18

MTCYBP23

MTND1P23

MTND2P28

MTND4P12

MTND5P11

NAP1L1P1

NBEAP1

NDUFAF4P3

NT5C3AP1

P2RX6P

PCBP2P1

PGAM1P8

PGBD4P1

PMP22

PPIAP22

PPIAP30

PPIAP31

PTMAP4

PTP4A2P2

RAC1P2

RIMKLBP1

RIMKLBP2

RN7SKP50

RNA5SP165

RNA5SP344

RNA5SP37

RPL23AP7

RPL23AP82

RPL31P7

RPL35P1

RPL37AP1

RPL7AP11

RPS15AP5

RPS7P3

RPSAP12

RRM2P3

SEC24AP1

SEPHS1P4

SHLD2P1

SMPD4P1

SUGT1P1

SUMO2P1

TAGLN2P1

TATDN2P1

TDGF1P5

TLK1P1

TMSB4XP1

TMSB4XP2

TMSB4XP4

TMSB4XP8

TPTEP1

TUBB3P2

UBE2V1P1

UPK3BP1

WHAMMP2

WHAMMP3

ZDHHC4P1
